# Scale-free models of chromosome structure, dynamics, and mechanics

**DOI:** 10.1101/2023.04.14.536939

**Authors:** Simon Grosse-Holz, Leonid Mirny, Antoine Coulon

## Abstract

The nucleus of a cell contains its genetic information in the form of chromatin: polymers of DNA and associated proteins. The physical nature of this polymer system is yet to be understood. Orthogonal experimental approaches probing chromosome structure, dynamics, and mechanics typically suggest the existence of scaling relationships, leading to the widespread use of scale-free, or fractal, models to represent interphase chromosomes. However, currently, there is no single physical model consistent with all reported scaling exponents. Here, we consider the space of possible scale-free models of chromosome structure, dynamics, and mechanics, and examine the fundamental connections between these physical properties. We demonstrate the existence of two algebraic relationships between the scaling exponents—–connecting structure with dynamics, and dynamics with mechanics, respectively–—outlining the necessary physical conditions for a model to match specific exponent values. Applied to values reported in metazoans, our theory identifies the family of models consistent with all observed scalings, which notably excludes the classical Rouse, Zimm, and fractal globule polymer models. Our theory highlights dynamic correlations between distal genomic loci as necessary to reconnect seemingly contradictory measurements. Consequently, we propose new experiments to narrow down the space of possible models. We expect this framework to serve as a guide for understanding past and future measurements, and for building new physical models of interphase chromosomes.

## Introduction

Chromosomes, in addition to being the carriers of genetic information, are long physical polymers made of DNA and associated proteins. Understanding their physical nature and spatiotemporal organization in the nucleus of a eukariotic cell is of central interest in the field. Specifically, the local chromosomal context of a given locus plays a crucial role in determining how the information encoded on the DNA is processed [1]. So-called *enhancer* elements for example are thought to activate their target genes by “looping in” and physically contacting the target promoter to initiate transcription [2–5]. How this interaction is regulated between elements that can be separated by millions of base pairs remains an open question [6–8]; in fact, even the structure, dynamics, and mechanics of the chromatin polymer itself—without reference to specific elements like enhancers and promoters—remain topics of active research [9– 12].

Our understanding of the 3D structure of interphase chromosomes has increased dramatically over the last decade, primarily due to experimental techniques like Hi-C [13, 14] (measuring pairwise contacts across the genome) and multipoint FISH methods [15, 16] (visualizing chromosome conformations in 3D space). Both techniques often report scaling relationships in a broad range of scales and organisms [9, 16– 21]. In metazoans specifically, exponents reported by both Hi-C and microscopy, from ∼10 kb to ∼100 Mb, suggest a *space-filling* organization of chromatin: two loci at a genomic separation *s* are on average separated in space by a distance 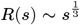 [9, 17], corresponding to a confining volume *V*(*s*) ∼ *R*^3^(*s*) ∼ *s*—thus the term “space-filling”. The probability *P*(*s*) of finding these two loci in contact is then given by the mean field approximation *P*(*s*) ∼ 1*/V* (*s*) [22]; *P*(*s*) ∼ *s*^−1^ was broadly observed in Hi-C and micro-C experiments across metazoans [13, 14, 19, 23]. Notably, this spacefilling spatial organization is more compact than one would expect for an ideal chain in equilibrium, which should adopt a random walk conformation with 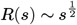, corresponding to 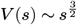 and 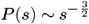 [24]; the latter, in turn, seems to be consistent with the situation in yeast [25].

Chromosome dynamics are usually studied by fluorescence microscopy, where current methods allow monitoring a few specific genomic loci, by targeting fluorophores either to exogenous DNA elements integrated into chromosomes [5, 26– 30] or by dCas9-based approaches [31–36]; alternatively, non-specific approaches such as labels on histone H2B can track many loci simultaneously, at the expense of knowledge or reproducibility of their genomic identity [10, 37–40]. With either of these imaging approaches ultimately measuring trajectories *x*(*t*) for genomic loci, quantification for these experiments usually employs the Mean Squared Displacement (MSD)

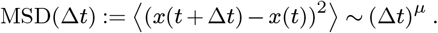

While a freely diffusive particle would exhibit a linear MSD curve (*µ* = 1), a chromosomal locus (i.e. point on a long polymer) is expected to move subdiffusively (*µ <* 1) due to the chain connectivity. Indeed, experiments show *µ* ≈ 0.5 − 0.6 in eukaryotic cells [25, 29, 30]. These values are consistent with the Rouse model of polymer dynamics, which predicts 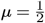 [41, 42]; which, however, also produces an ideal chain structure, 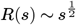, inconsistent with experimental data.

Taking an orthogonal angle on the question of chromatin dynamics, the present authors, together with others, recently presented an experimental system to measure the force response of a single genomic locus [43]. In response to a constant force switched on at *t* = 0 the locus moved as *x*(*t* ; *f*) ∼ *t*^0.5^, consistent with the same (Rouse) model for polymer dynamics that predicted the MSD scaling *µ* = 0.5—but which is inconsistent with the structure 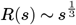 of real chromatin.

A model consistent with the space-filling structure of metazoan chromatin is the *fractal globule*, which describes crumpling of the chain due to topological constraints [44]. This, however, predicts an MSD scaling of 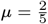 [45], markedly lower than the *µ* ≈ 0.5 − 0.6 observed in experiments.

In light of these observations, neither Rouse nor the fractal globule can provide satisfactory models for chromosome organization; which should not be surprising, given that these simple toy models neglect almost all biological features like loops, TADs, compartments, and active dynamics in the nucleus. More detailed models, usually computational, are thus being used by the community to investigate these biological features and their interplay. Notably, however, reference to scaling exponents—like *µ* and *ν* above—is still frequently made in discussing these results, implying that scale-dependent features like TADs are seen as “added on top of” some scale-free background model [46, 47]. This mindset is supported by the experimental observation that removal of some of these specific features, e.g. loops, broadens the region where *P* (*s*) can be well approximated by scaling relationships [19].

The main purpose of the present work is to point out that currently we do not have such a scale-free background model that would be consistent with the reported scaling exponents. While chromosome structure looks fractal-globule-like, dynamics and mechanical responses look Rouse-like; there is no understanding as to how to fit these observations together. We propose two avenues towards resolving this issue: first, we can question the usefulness of simplistic toy models. Specifically, one could argue that discussing scaling exponents for chromosome organization is quite meaningless, since it is dominated by scale-dependent biological processes and simply not well approximated by scale-free models that neglect these details. In this understanding, powerlaw scalings and associated exponents reported from experimental observations would fall prey to the quip known as Mar’s law (see also [48]): “everything is linear if plotted log-log with a fat magic marker.” Biological observables are never truly power-laws and thus do not exhibit scaling exponents. The alternative to this radically sober approach is to take the scaling approximation seriously: assuming that experimental observations are indeed reasonably well approximated by powerlaws, we can attempt to construct a scale-free model consistent with the reported exponents, to serve as rudimentary approximation to chromosome organization.

The present work pursues this second line of argument: assuming that it is in fact a useful approximation, what are the implications of a consistent, scale-free model of chromosome organization?

## Results

Let us construct such a scale-free model. By “scale-free”, we mean that there is no intrinsic length scale in the system; so we idealize a chromosome as an infinitely long polymer (no large length scale) without any microstructure such as monomer size (no short length scale). Any such model should make predictions for the observables discussed in the introduction; and because of the absence of finite scales, we expect to find powerlaws. Explicitly, we assume the forms

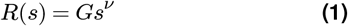

for the root-mean-square spatial distance between two loci at a genomic separation *s*;

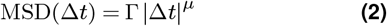

for the MSD of a single genomic locus; and

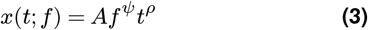

for displacement of a single locus in response to a constant force *f* switched on at time *t* = 0 (red, teal, and orange in fig. 1).

**Figure 1.**
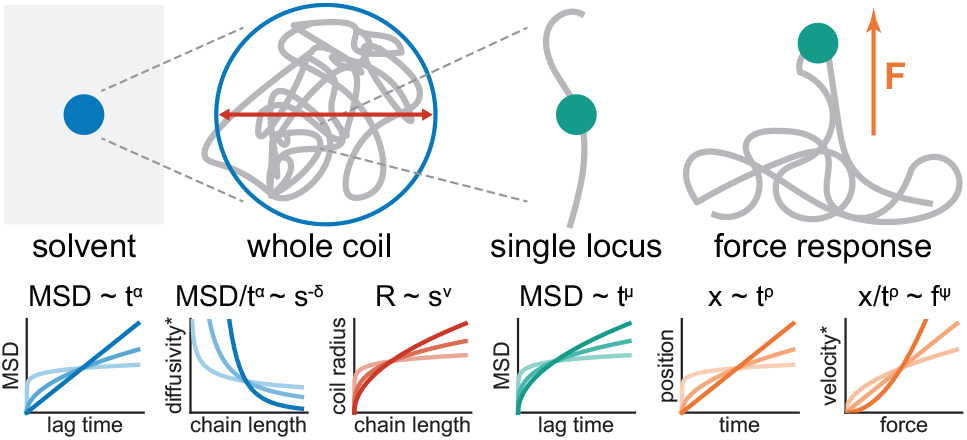
Summary of the exponents considered in the text and what part of the system they relate to. Left to right: *α* describes the viscoelasticity of the solvent, i.e. the MSD of a free tracer particle. Considering an isolated polymer coil as such a tracer particle, *δ* reflects the dependence of its (anomalous) diffusivity on the chain length; *ν* gives the scaling of the physical radius of the coil. The motion of individual loci within the coil is characterized by *µ*. Upon application of an external force, such loci exhibit a powerlaw response with exponent *ρ*; the (fractional) velocity of this response is force dependent, with exponent *ψ*. Colors indicate which constitutive relation an exponent is associated with: red for eq. (1), teal for eq. (2), orange for eq. (3), and blue for eq. (4). *: “anomalous diffusivity” if *α* ≠ 1 “fractional velocity” if *ρ* ≠ 1.

While the last two observables (dynamics and force response) probe individual loci, the first one (structure) probes finite subchains. To bridge this gap, we consider the whole-coil diffusion of a finite and isolated subchain (i.e. we disconnect it from the infinite polymer and place it in empty solvent). Over timescales longer than the internal relaxation time of this coil, we expect it to diffuse in the solvent like a free particle, with a diffusion constant dependent on its size *s*, i.e. an MSD of the form

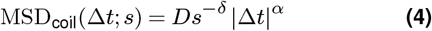

(blue in fig. 1). Since we expect a free coil/particle to undergo normal diffusion, *α* = 1 seems like the most natural choice; however, allowing *α <* 1 incorporates the possibility of a viscoelastic solvent, such that even a free tracer particle would undergo subdiffusion—which has been observed for the nucleoplasm, though estimates for *α* vary broadly (*α* ≈ 0.5 − 1) [49–51]. The exponent *δ* can be understood as incorporating dynamic correlations of different loci along the polymer: for a freely draining chain (such as the Rouse model), monomers are independent from each other and whole-coil diffusivity is simply inversely proportional to chain length, yielding *δ* = 1. The Zimm model [42], in contrast, incorporates hydrodynamic interactions between the loci, which results in a hydrodynamic radius 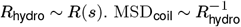 then implies *δ* = *ν*.

Through eqs. (1) to (4), our scale-free description of chromosome organization is characterized by the model constants *G*, Γ, *A, D*, and an energy scale *k*_B_*T* ; which have dimensions

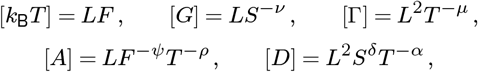

where we use the symbols *L, F*, *S, T* to denote length, force, genomic distance, and time, respectively. We can combine these model constants into new quantities

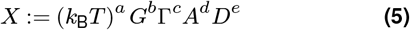

with units

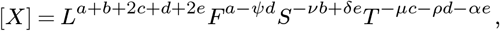

which allows us to construct e.g. a length scale by solving a simple linear system: we set [*X*] = *L*, which gives a system of four equations for the five variables *a, b, c, d, e*; elementary substitutions reduce this system to one equation for two variables:

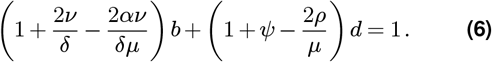

This equation has a one-parameter family of solutions (*b, d*), so long as either of the terms in brackets is non-zero. We have thus constructed a length scale *X* from the model constants of our—supposedly—scale-free approximation, showing that said approximation is inconsistent; unless the model exponents satisfy

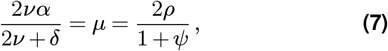

ensuring that both brackets in eq. (6) vanish. These exponent relations are therefore required for self-consistency of the scale-free approximation: if they are violated, we can explicitly construct a length scale from the model constants.

Our derivation of eq. (7) relies only on dimensional analysis, emphasizing that these relations are a direct consequence of the scale-free assumption, i.e. the same assumption that lets us approximate individual observables as powerlaws in the first place. The same relations can, however, be obtained by other arguments: in appendix A we demonstrate the physically more intuitive approach *via* crossover points between the different scaling regimes of a locus in a finite coil; in appendix B we discuss how eq. (4) (free diffusion of a finite coil) can be replaced by other observables with the same dimensional structure (depending on a time lag Δ*t* and polymer separation Δ*s*); finally, appendix C gives an explicit formulation in terms of the polymer configuration *x*(*s, t*). Special cases of these arguments exist in the literature and specifically the first relation in eq. (7) has been reported previously: in the cases *α* = 1, *δ* = 1 [45, 52]; 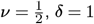 [53]; and *δ* = 1 [54]. The second relation connecting dynamics and force-response is satisfied explicitly by the Rouse model [43, 55], but has not been studied in generality.

## Discussion

Having established eq. (7) as the central consistency condition for the scale-free approximation in chromosome organization, we now discuss its implications in light of available experimental evidence (table 1). We find that existing observations make specific predictions for less commonly studied exponents and we propose experimental approaches to measure those.

**Table 1.** Measured scalings for *R*(*s*) ∼ *s*^*ν*^, MSD(Δ*t*) ∼ (Δ*t*)^*µ*^, and *x*(*t*; *f*) ∼ *t*^*ρ*^. Gray background highlights reporting of multiple exponents from the same experimental system. Chromosome structure is frequently not strictly fractal due to loop extrusion; therefore we here focus on experiments where loop extruding factors (RAD21, a component of the cohesin complex) or their loaders (NIPBL) were acutely degraded, where possible. This overview is not exhaustive.

| Organism | $\nu$ | $\mu$ | $\rho$ | Ref. | Notes |
| --- | --- | --- | --- | --- | --- |
| *H. sapiens* |  |  |  |  |  |
| HCT-116 | 0.3-0.4<sup>1</sup> | - | - | [18] | ΔRAD21 |
|  | 0.3-0.4<sup>2</sup> | - | - | [16] |  |
| HeLa | - | 0.5 | - | [56] | Telomeric probes |
| U2OS | - | 0.5 | 0.5 | [43] |  |
|  | - | 0.55 | - | [56] | Telomeric probes |
| MF | - | 0.7 | - | [56] | Telomeric probes |
| *M. musculus* |  |  |  |  |  |
| mESC | - | 0.5<sup>3</sup> | - | [29] | WT and ΔRAD21 |
| | - | 0.6<sup>3</sup> | - | [30] | WT and ΔRAD21 |
| | 0.33-0.4<sup>1</sup> | - | - | [21] | ΔRAD21 |
|  | 0.15-0.4<sup>2</sup> | - | - | [9] |  |
| hepatocytes | 0.4<sup>1</sup> | - | - | [19] | ΔNIPBL |
| 3T3 | - | 0.4 | - | [56] | Telomeric probes |
| *D. melanogaster* | 0.31<sup>3</sup> | 0.52<sup>3</sup> | - | [5] |  |
|  | 0.22-0.37<sup>2</sup> | - | - | [17] |  |
| *S. cerevisiae* | 0.5<sup>1</sup> | - | - | [20] |  |
|  | - | 0.5 | - | [57] |  |
| *E. coli* | - | 0.4 | - | [58] |  |
| *Caulobacter* | - | 0.4 | - | [58] |  |

<sup>1</sup>from Hi-C contact probability $P(s) \sim s^{-3\nu}$ [22]
<sup>2</sup>direct measurement from multiplexed FISH
<sup>3</sup>two-locus live-cell measurement

### Mechanics & Dynamics

Consider the force response experiments of [43], where we determined, in the same system, *ρ* ≈ 0.5, *ψ* ≈ 1, and *µ* ≈ 0.5, fully consistent with eq. (7). Notably, just the linear force response (*ψ* = 1) suffices to predict *ρ* = *µ*; our measurement of the force response exponent *ρ* ≈ 0.5 can thus be interpreted as an independent validation of earlier experiments finding *µ* ≈ 0.5 (table 1). In addition, our theory predicts that if *ρ* ≠ *µ* (e.g. future experiments in different chromatin contexts, different cell types, or different spatial, temporal, or force regimes), this would hint at a non-linear force response *ψ* ≠ 1.

### Structure & Dynamics

The first relation in eq. (7) connects the structural and dynamical scalings *ν* and *µ*, both of which have been investigated in various experimental systems (see table 1). While specifically yeast seems consistent with the expectations for a Rouse model, i.e. *µ* = 0.5, *ν* = 0.5, and *α* = 1 [59], metazoans like fruit fly, mouse, or human, seem to behave differently. For the purpose of this discussion, let us consider the case *µ* = 0.5, *ν* = 0.33 (fig. 2); this seems consistent with best estimates, but is of course an idealization of the experimental situation. While we choose this idealization for consistency with the literature models mentioned in the introduction (Rouse: *µ* = 0.5; fractal globule: *ν* = 0.33), we emphasize that eq. (7) holds for any value of these exponents and similar discussion can be furnished with different numerical values.

**Figure 2.**
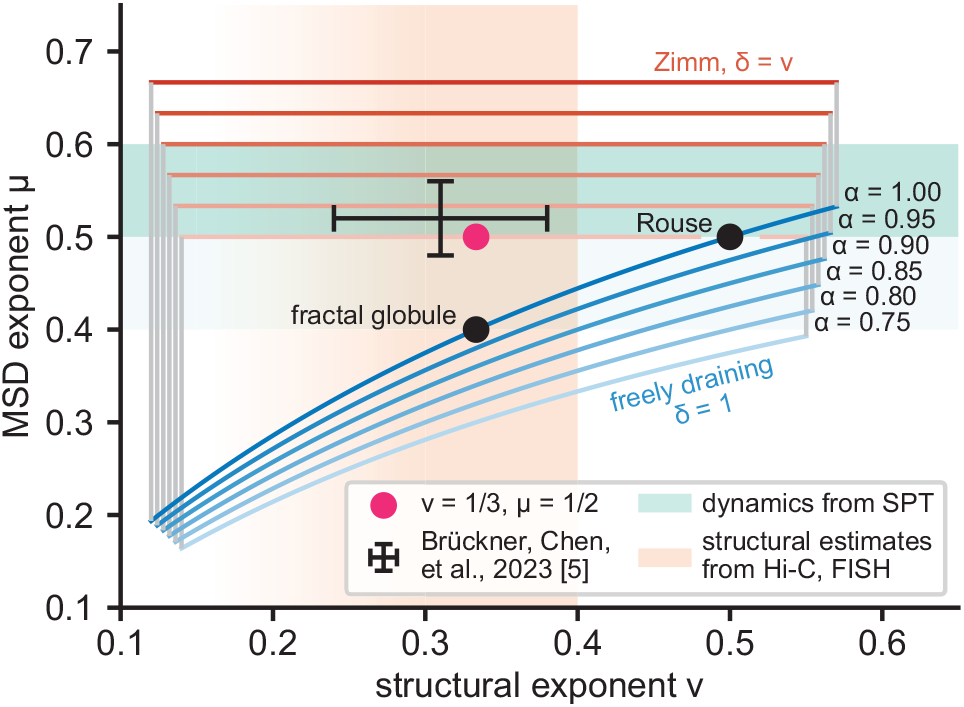
Experimental results in the context of eq. (7). Shaded regions are consistent with experimental determinations of structure *ν* (orange) or dynamics *µ* (green, from SPT; dense shade indicates eukaryotic estimate *µ* ≈ 0.5 − 0.6, light shade extends to bacterial estimate *µ* ≈ 0.4) respectively. Black error bars indicate estimate from [5]. Red circle marks *ν* = 0.33, *µ* = 0.5, which serves as example for discussion in the main text. Outlines show theoretically plausible regions (eq. (7)) for different *α*, as indicated. The top (red) edge of these regions is given by the Zimm condition *δ* = *ν*, while the bottom (blue) edge is given by the freely draining chain (*δ* = 1); the points inbetween correspond to *ν < δ <* 1. Horizontal cutoffs (gray lines) are chosen for visual appeal. Common polymer models: Rouse chain and fractal globule are indicated as black circles; both are instances of a freely draining chain with *α* = 1 (blue curve).

Reformulating the first relation in eq. (7) as

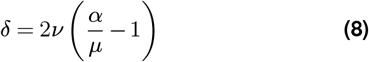

shows that we should expect a 1-parameter family of models with different *α* and *δ* that exhibit the desired scalings in *µ* and *ν*. We discuss a few of these options:

- In a freely draining chain (blue lines in fig. 2), individual monomers are independent, such that *δ* = 1. This assumption is made in the Rouse model and in [45] for dynamics of the fractal globule. Equation (8) then becomes 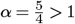, i.e. we would need a medium in which free tracers undergo *superdiffusion*. This appears unrealistic for the nucleoplasm. While superdiffusion of free tracers might theoretically be achieved by energy dependent processes like transcription or loop extrusion, those are unlikely to be scale-free, such that we will not further pursue this point here.
- Zimm’s treatment of hydrodynamic interactions between different monomers amounts to *δ* = *ν*, such that 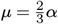 independent of *ν* (red lines in fig. 2). This would allow matching *µ* ≈ 0.5 by tuning *α* ≈ 0.75. While this is within current estimates for nucleoplasm viscoelasticity, these estimates scatter quite broadly (*α* ≈ 0.5 − 1), such that this consistency statement is rather weak. Furthermore, due to crowding one might expect hydrodynamic interactions to be screened in the nucleus [60, 61], such that *δ* = *ν* appears questionable in the first place.
- Between the two canonical values of *δ* = 1 (freely draining chain) and *δ* = *ν* (Zimm-style hydrodynamic interactions), it is conceivable that chromatin loci in the nucleus exhibit dynamic correlations with *ν < δ <* 1. In a purely viscous nucleoplasm (*α* = 1), eq. (8) would then imply 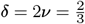, i.e. a whole-coil hydrodynamic radius scaling with the crosssection of the coil: *R*_hydro_ ∼ *R*^2^(*s*).
- We are not aware of a physical model that would produce this intermediate level of dynamic correlations between different loci. It seems conceivable, however, that hydro-dynamics plays a role. The Zimm model (cf. previous point) treats hydrodynamics as an effective two-body interaction, which might overestimate the correlations created in a densely packed chromosome: other parts of the chain might be close enough to screen hydrodynamic interactions between two loci—thus increasing *δ > ν*— without going so far as to annihilate the effect completely. It remains to be seen in future studies whether this provides a viable explanation. At the whole-nucleus scale, correlated dynamics have been observed [11] and attributed to hydrodynamic interactions [62].

We note that eq. (8) allows us to reconcile any arbitrary combination of exponents (*ν, µ*) (such as our discussion example (0.33, 0.5)) by hypothesizing suitable values for *δ*, which has received little experimental attention to date. But eq. (8) is an experimentally testable prediction and we propose to test it with experiments discussed in the next section.

### Proposed experiments

The self-consistency of the scale-free approximation is experimentally testable. There are various experimental systems that might be employed (fig. 3), which we discuss in the following.

**Figure 3.**
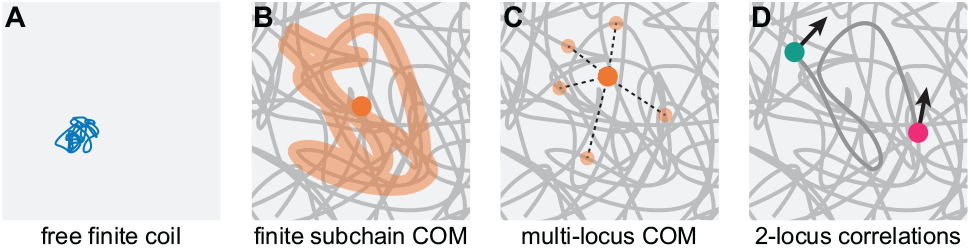
Overview of proposed experimental approaches to completing the scale-free null model. Left to right: diffusion of a free coil in otherwise pure solvent over long time scales; diffusion of the center of mass of a finite subchain at short times; diffusion of the center of mass of multiple tracer loci at short times; displacement correlation/covariance of two tracers with known genomic separation at short times. These experimental setups correspond to observables characterized by (*α, δ*) (main text), (*α*^*′*^, *δ*^*′*^) (appendix B), eq. (23), and (*α*^*′′*^, *δ*^*′′*^) (appendix C), respectively. See discussion in the main text.

- *Free, finite coil* (fig. 3A). Equation (4) describes the free diffusion (in solution) of finite stretches of chromatin of different lengths. Possible experimental realizations are micro-injected nucleosome arrays [63], extrachromosomal DNA [64], or chromatinized DNA fragments in nuclear extract. We note that in order to measure the exponent *δ*, these experiments would have to be run for a range of chain lengths *s*. Furthermore, a major issue for these approaches would be to verify that the structure of the chromatin fragments still obeys eq. (1). Finally, what exactly constitutes the “solvent” in these scenarios can vary: while the picture presented here assumes that the tracked fragment moves in a chromatin-free environment (e.g. nuclear extract), for e.g. extrachromosomal DNA one would abstract the surrounding chromatin in the nucleus into an effective solvent; the measured exponents (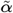 and 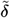 in appendix B) would be different from *α* and *δ* here, but still satisfy a relation of the form eq. (7). See eq. (10) and pertinent discussion. The advantage of using eq. (4) is that the exponent *α* describes the free (sub-)diffusion of a generic tracer particle in the solvent; it is thus often interpreted as a property of the medium alone and of interest in its own right.
- *Center of mass of a finite subchain* (fig. 3B). In appendix B we argue that short-time dynamics of the center of mass of a finite subchain—which is still connected to the infinite polymer, as opposed to the free coil discussed before—obeys an equation (10) that is analogous to eq. (7), albeit with different exponents (*α′, δ ′*). However, even though techniques for fluorescent genome labeling have recently enabled chromosome tracing in fixed cells [9, 15–17], labeling a continuous genomic region in live cells seems difficult with current techniques and has not been reported yet. So while eq. (10) does provide an alternative to eq. (7) that does not require excision of a part of the chain, it still seems out of reach of current experiments.
- *Center of mass of multiple loci* (fig. 3C). While homogeneous labeling and tracking of extended genomic regions in live cells is currently not possible, tracking multiple point-like loci with defined genomic identities has been demonstrated with various methods [5, 26–30, 32–36]. The center of mass of such a multi-point system exhibits dynamics that are non-trivial; specifically we expect an intermediate regime where the MSD is not necessarily a powerlaw, due to the abundance of length scales introduced by the label positions. The underlying chromosome organization, however, can still be approximated as scale-free, which now leads to a self-consistency condition in the form of eq. (23). Such multi-point experiments are feasible with current technology, but existing data sets do not suffice to validate our predictions, since our framework requires making MSD measurements on the same system for loci arranged across regions of different genomic sizes. In addition, issues with existing data approaching these requirements [5, 35] are a low number of tracked loci, low temporal resolution, and global nuclear movement. We note that since this approach relies only on the center of mass of multiple loci, it is not necessary to be able to distinguish the different loci or track them individually; they might thus be labelled in the same color, which drastically increases the number of loci that can be imaged simultaneously.
- *Two-particle displacement covariance* (fig. 3D). For short lag times Δ*t*, the instantaneous covariance of displacements of two loci on the chain follows eq. (21) and the pertinent exponents (*α*′′, *δ*′′) satisfy the same relation as (*α, δ*) for the free coil. Two-locus tracking experiments have been performed recently [5, 26–30], although out of those only [5] was able to vary the genomic distance between the loci, which is necessary to measure *δ*′′. Repeating their experiments at higher time resolution should make these exponents accessible.

## Conclusion

Scaling behavior of different aspects of chromosome organization is reported widely in the literature. We have demonstrated that self-consistency of the underlying approximation—that of the absence of relevant length scales—establishes non-trivial connections between the pertinent scaling exponents. These relations highlight that dynamic correlations between different genomic loci are the missing piece to the puzzle of reconciling chromosome structure, dynamics, and mechanics. We close by discussing possible experiments to address this gap.

## Acknowledgements

We are grateful for productive discussions with many colleagues, including Vittore Scolari, Kirill Polovnikov, and members of the Mirny and Coulon groups; Mikhail Tamm; Jean-François Joanny; and Alexander Grosberg. Françoise Brochart-Wyart and her work provided valuable inspiration. This work received funding from the Centre National de la Recherche Scientifique (CNRS), the Institut Curie, the European Research Council (ERC) under the European Union’s Horizon 2020 research and innovation program (grant agreement 757956), the LabEx CELL(N)SCALE and the LabEx DEEP (ANR-11-LABX-0038, ANR-11-LABX-0044 and ANR-10-IDEX-0001-02), the Agence Nationale de la Recherche (project CHROMAG, ANR-18-CE12-0023-01; project MechaChrom ANR-22-CE95-0003-01), the National Science Foundation (NSF-ANR: Physics of chromosomes through mechanical perturbations, NSF 2210558), the MIT-France Seed Fund, and the Chaire Blaise Pascal program of Île-de-France. Our figures use Paul Tol’s *Vibrant* color scheme [65]; this PDF uses the LATEX template found at github.com/quantixed/manuscript-templates.

## APPENDIX

### A. Finite subchain

The argument in the main text is formulated in terms of dimensional analysis to emphasize that it is a necessary conclusion of the scale-free assumption. It is easily reformulated in a more physical language by considering a finite subchain of length 𝔰 [54].

Equation (1) gives the physical size of this subchain as

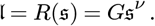

This allows for two independent definitions of a time scale: by setting MSD(Δ*t*) = 𝔩 ^2^ we find

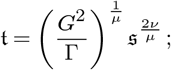

letting instead MSD_coil_(Δ*t*; *s*) = 𝔩 ^2^ yields

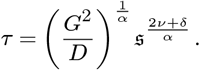

Physically, both describe the relaxation time scale of the coil and should thus be equal up to numerical prefactors. This requires that the exponents on 𝔰 be the same, yielding the first relation in eq. (7).

Similarly, eqs. (2) and (3) and the thermal energy *k*_B_*T* allow for the construction of two force scales associated with our subchain, both of which should exhibit the same scaling behavior with 𝔰 (or in this case 𝔩):

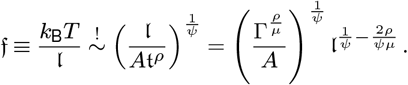

Again equating the exponents (on 𝔩) yields the second relation in eq. (7).

While the formulation in terms of a finite subchain can aid physical intuition, the core argument remains the same: if eq. (7) does not hold, a finite length scale emerges. To see this, consider:

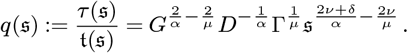

Since *q* is a dimensionless ratio, if the scaling with s were non-trivial, the combination of constants in front would have units of *S* to some power, translating to a length scale through eq. (1). Thus, within the framework of scale-free models, any finite scale (length, force, or otherwise) associated with the subchain s has to be unique. This is ultimately what drives the scaling argument.

### B. Alternatives to eq. (4)

The argument in the main text relies on the long-time asymptote of free coil diffusion to make the link between structure and dynamics. We chose this approach because it directly connects to the diffusion of free tracer particles in the solvent, which has previously been investigated experimentally. Its downside, however, lies in having to “cut out” a piece from the infinite system; we are thus technically considering two distinct systems, which might create conceptual confusion. Here we demonstrate that the same results hold for other observables that have the same dimensional structure as eq. (4), without the need to cut a piece from the chain.

Consider the center of mass of a finite subchain of length *s*. Over times much longer than the internal relaxation of the sub-chain, the subchain moves as a whole; the motion is then dominated by its connection to the infinite chain and the center of mass MSD asymptotes to the MSD of a single locus,

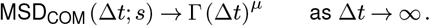

The short time behavior of the center of mass, however, introduces a new law

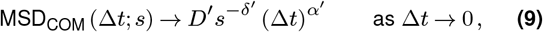

with exponents *δ*′ and *α*′ which are *a priori* undetermined and unrelated to *δ* and *α* of the main text. After substituting eq. (9) for eq. (4), the dimensional analysis follows exactly the same lines as in the main text, yielding the equivalent of eq. (7):

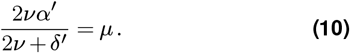

So short-time center of mass diffusion of a finite subchain satisfies the same exponent relation as long-time diffusion of a free, finite coil; this relation is forced by the scale-free assumption in either case. Note that this does not imply that *α*′ = *α* and/or *δ*′ = *δ*; even though especially the latter seems plausible on physical grounds, it is not required by the scale-free assumption itself.

Similarly, we might consider the correlated displacement of two loci *s* and *s*′ on the chain (see also eq. (21)). Over lag times shorter than the equilibration time of the tether between the loci, the covariance of their displacements is expected to follow powerlaws in lag time and tether length, with exponents that we can call *α*′′ and −*δ*′′ respectively. Again, dimensional analysis follows the same lines as before and we find a relation of the form eq. (10), with double-primed exponents substituted for single-primed ones.

Clearly, any observable that dimensionally matches eq. (4) (i.e. is a powerlaw in both genomic separation and lag time) has to satisfy the appropriate version of eq. (7). This argument allows us to extend the discussion of the long-time free coil diffusion of the main text from the behavior in empty solvent (as we assumed initially) to solvent that still contains the infinite “rest” of the chain (which might be experimentally more realizable). At long times we still expect diffusion of the form (4), however now with independent exponents 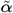 and 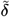, satisfying eq. (10) (upon replacing primed with tilde exponents). The exponent 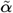 can now be interpreted as characterizing the background polymer solution, where *α* in the main text characterizes the solvent itself without polymer.

### C. Explicit polymer

In the main text we emphasized that the presented exponent relations follow directly from the scale-free assumption by simple dimensional analysis. It is also possible to phrase the same ideas as an explicit critical (i.e. scale-free) field theory for the polymer conformation *x*(*s, t*); *s* being the coordinate along the polymer backbone and *t* time.

We express the two-point correlations of the conformation *x*(*s, t*) as

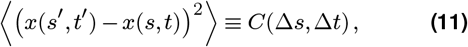

where here and in the following we use the shorthand Δ*s* ≡ *s*′ − *s* and Δ*t* ≡ *t*′ − *t*. We assume parity symmetry (i.e. invariance under *s* → −*s*), such that *C*(Δ*s*, Δ*t*) = *C*(|Δ*s*|, Δ*t*); by symmetry of the expression on the left hand side of eq. (11) we then have *C*(Δ*s*, Δ*t*) = *C*(|Δ*s*|,|Δ*t*|). We assume that the system is scale-free, i.e. there exist exponents *a, b*, and *c* such that for any *r >* 0

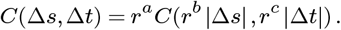

Formally, we can then set *r* = |Δ*t*|^−1*/c*^, yielding

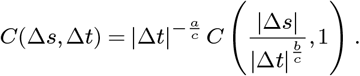

We rename the exponents 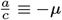 and 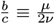, introduce constants Γ and 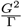, and define

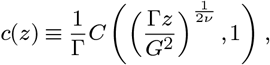

such that ultimately we can write the generic form of the two-point correlations in our scale-free model as

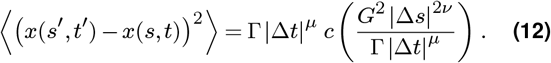

This provides the starting point for our discussion.

For eq. (12) to describe a meaningful polymer model, we require that the limits Δ*t* → 0 (structural scaling) and Δ*s* → 0 (single locus MSD) are finite; with appropriate choice of the constants Γ and 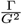 this then enforces

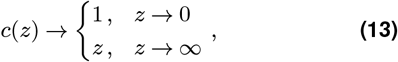

such that the two limits reproduce eqs. (1) and (2) of the main text, respectively. Convergence for *z* → ∞ is understood as 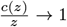 here, though we show below that some weak assumptions for “physical” polymer models are equivalent to enforcing the stronger *c*(*z*) − *z* → 0 (which we would write as *κ* ≥ 0^+^ in the notation below).

Consider now a linear observable

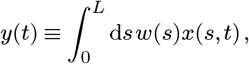

where *L* is a constant (length along the polymer) and *w*(*s*) is some weight distribution with supp *w* ⊆ [0, *L*]. Examples include a single locus, *w*(*s*) = *δ*(*s*); the relative position of two loci, *w*(*s*) = *δ*(*s* − *L*) − *δ*(*s*); or the center of mass of a finite subchain, 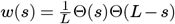; where *δ*(*s*) is Kronecker’s *δ*-distribution and Θ(*s*) the Heaviside-Θ function.

We can write the MSD of the observable *y*(*t*) as

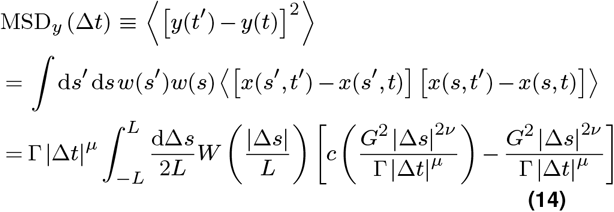

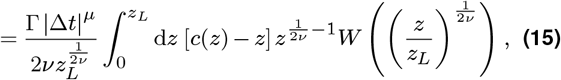

where the first step is the definition of MSD and the second inserts the definition of *y*(*t*); in the third step we expand the terms inside the expectation value and complete the squares to produce square terms for which we can then substitute eq. (12); at the same time, we introduce

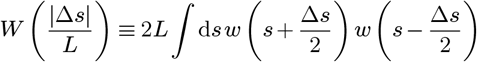

to reparametrize the double integral; in the last step we exploit the symmetry in the integrand to restrict the integration domain to the positive half axis and substitute integration variable and domain boundary

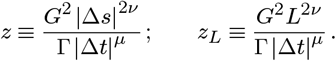

Note that if *W* (*a*) has finite weight at *a* = 0 (i.e. a component *δ*(*a*)), the lower limit of the integral in eq. (15) should be understood to give half of that contribution for the result to be consistent with eq. (14).

#### Limit of long lag times

Consider the long-time limit Γ|Δ*t*|^*μ*^ ≫ *G*^2^*L*^2*ν*^. Then 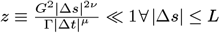 and consequently *c*(*z*) → 1 uniformly over the whole integration domain in eq. (14). Introducing the constants

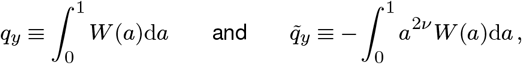

we then find

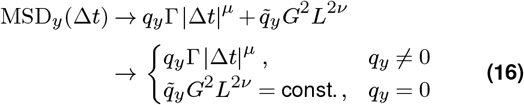

as Δ*t* → ∞.

The numerical prefactors *q*_*y*_ and 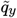 depend on the construction of the observable *y*(*t*): for a single locus (*w*(*s*) = *δ*(*s*)) one finds 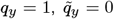; center of mass of a subchain of length *L* gives 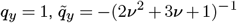; while for the relative position of two loci (*w*(*s*) = *δ*(*L* − *s*) − *δ*(*s*)) one finds 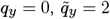. Note the limiting behavior for *q*_*y*_ = 1, which are “center-of-mass-like” observables: over long times, those behave like single loci,

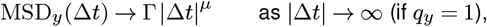

independent of their internal makeup.

#### Two-point MSD

We consider the relative motion of two loci at a fixed separation *L*; interestingly, this observable turns out to provide a sufficient characterization of the whole model, at the level of description employed here (two-point correlations).

Let *w*(*s*) = *δ*(*L* − *s*) − *δ*(*s*); then

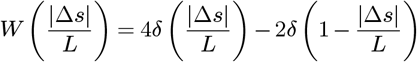

and eq. (14) becomes

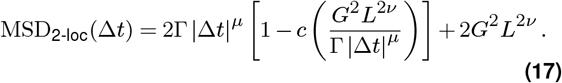

Note that with the asymptotics of *c* given in eq. (13), this produces the expected limits

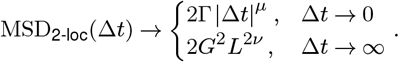

Thus, knowing MSD_2-loc_ for multiple pairs of loci at different separations *L* (as in e.g. [5]) allows us to identify *µ, ν*, Γ, and *G* from the asymptotes and then *c*(*z*) from the behavior at finite lag times. The two locus MSD is thus a sufficient characterization of the whole scale-free model; we exploit this statement in an example below to indentify *c*(*z*) for the continuous Rouse chain.

#### Center of mass of a finite subchain

As pointed out in a separate section of this appendix, the dimensional treatment for this observable is exactly the same as for the finite coil in the main text, with the difference that the exponent *α*′ now does not relate to the diffusion of free tracer particles and thus cannot be interpreted as an intrinsic property of the solvent. So here we simply expect to reproduce eq. (10).

Let 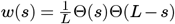. Then

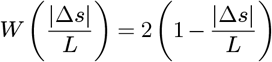

and eq. (15) becomes

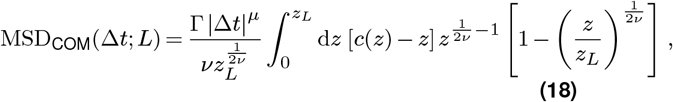

where we remind ourselves of the time dependence of

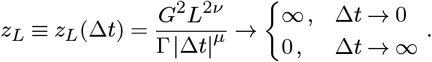

Taking the long time limit *z*_*L*_ → 0 of eq. (18) we note that *c*(*z*) − *z* → 1 ∀*z* ∈ [0, *z*_*L*_], such that we can substitute 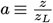 in the integral and obtain

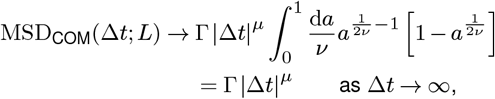

as expected from the more general treatment of eq. (16). For the short time limit *z*_*L*_ → ∞ we introduce

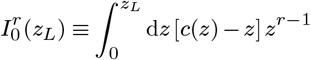

and rewrite the integral in eq. (18) as

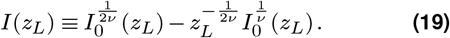

We introduce the asymptotic expansion

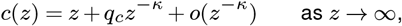

with *q*_*c*_ ≠ 0 and *κ >* − 1 (such that 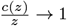 as *z* → ∞, as required by eq. (13)). For large *z*, the dominant term in the integrand of 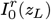 is then 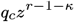, such that

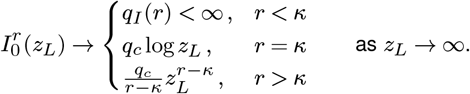

Applying this to the second term in eq. (19), we note that the prefactor kills all the cases where the integral does not diverge fast enough:

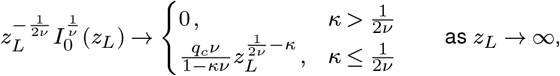

such that overall eq. (19) has the limiting behavior

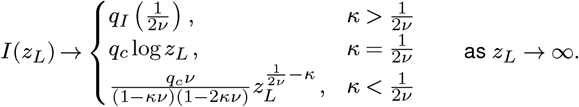

We summarize this behavior as

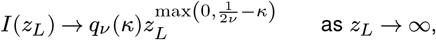

With

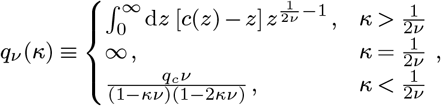

where 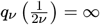 encodes the logarithmic correction and matches the limiting behavior from both sides.

Upon inserting into eq. (18), we define 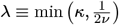 and find (as *z*_*L*_ → ∞)

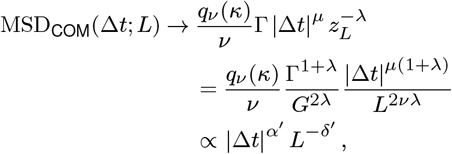

where in the last step we introduced *α*′≡ *µ*(1 + *λ*) and *δ*′≡ 2*νλ*, which are the heuristic exponents for short time center of mass diffusion that we used in eq. (10).

It is now straightforward to check that the parametrization of *α*′ and *δ*′ in terms of *λ* is equivalent to eq. (10): for any *λ* ∈ ℝ,

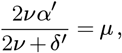

while starting from that equation we can rewrite it as 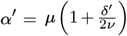 and introduce 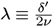 to reobtain *α*′ = *µ* (1 + *λ*) and *δ*′ = 2*νλ*. Ultimately, we have recovered from the explicit model the exponent relationship previously derived from dimensional analysis.

The explicit model does, however, allow us to make additional connections that are not clear from dimensional analysis, as demonstrated in the next section.

#### Displacement covariances

Consider two loci positioned at *s* and *s*′ along the chain and their displacements over a finite lag time Δ*t* ≡ *t*′ − *t*. These displacements have a covariance

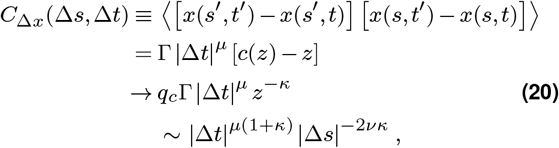

where the second step is analogous to eq. (14), we consider the limiting behavior for early times, Δ*t* → 0 (equivalently *z* → ∞), and use *q*_*c*_, *κ*, and *z* as defined previously. This short-time asymptote has the form

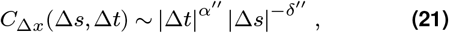

where the exponents *α*′′≡ *µ*(1 + *κ*) and *δ*′′≡ 2*νκ* satisfy the same exponent relationship as *α*′ and *δ*′ in eq. (10), or *α* And *δ* in eq. (7), as expected from the dimensional argument.

Note, however, that *δ*′ and *δ*′′ both rely on the same physical phenomenon: correlated motion of different loci at early times, quantified by *κ*; one finds

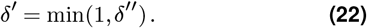

For an intuitive interpretation of this relation, note that *δ*′′ encodes how the movements of two loci on the chain—separated by a finite distance Δ*s*—are correlated at short times Δ*t*, i.e. before displacements can propagate along the backbone. That is, *δ*′′ encodes *non-backbone mediated correlated motion* of two loci. If these correlations are long-ranged (*δ*′′ *<* 1), they speed up the center of mass diffusion of finite coils (*δ*′ = *δ*′′); if they are short-ranged (*δ*′′ *>* 1), finite coil diffusion is governed by the backbone alone and *δ*′ = 1. Intuitively one would similarly expect *δ* = *δ*′, but this remains outside the realm of the present model, as *δ* in the main text characterizes the motion of a finite stretch of chain, not a subchain of the infinite system.

Another interesting observation concerning eq. (20) is that it allows us to further constrain the generic scale-free model. While so far we considered *κ >* −1 (which is enforced by the requirement that 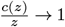 as *z* → ∞), we now note that *κ* ≤ 0^−^ would imply *δ*′′≤ 0^−^, i.e. short-time displacement covariance *increases* with increasing separation of the loci under study; which seems pathological for the polymer models we consider here. The limiting case *κ* = 0, in turn, is more interesting: in this case *δ*′′ = 0, i.e. short-time displacement covariance decays slower than any power with Δ*s*; or not at all. Specifically, they might be completely independent of Δ*s*, in which case

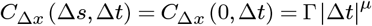

implies *c*(*z*) = *z* + 1 and specifically *q*_*c*_ = 1. This scenario describes a frozen polymer configuration moving as a rigid object and is probably the most intuitive instance of the edge case *κ* = 0.

#### Center of mass of multiple loci

Let 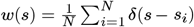, with *s*_*i*_ ∈ [0, *L*] ∀*i*. Then

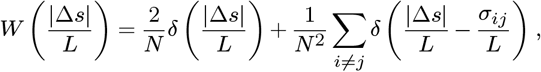

where *σ*_*ij*_ ≡ |*s*_*i*_ − *s*_*j*_ |, such that eq. (14) becomes

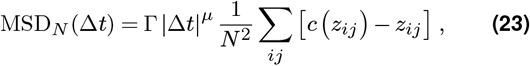

with

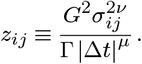

At long times *z*_*ij*_ →_*µ*_ 0 ∀*i, j*, such that *c*(*z*_*ij*_) → 1 and MSD_*N*_ (Δ*t*) → Γ|Δ*t*|^μ^ as expected by eq. (16).

At short times *z*_*ij*_ → ∞ ∀*i* = *j* while clearly still *z*_*ij*_ = 0∀*i* = *j*, such that

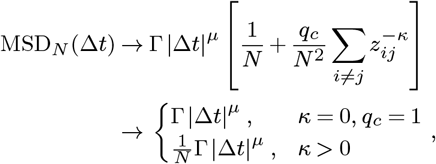

as Δ*t* → 0, where we omit the cases *κ <* 0 and *q*_*c*_ ≠ 1 if *κ* = 0, in line with the discussion of *κ* above. The case *κ* = 0, *q*_*c*_ = 1 corresponds to a frozen coil as discussed before, while *κ >* 0 matches the expectation that at short times different loci move independently, such that 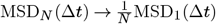. Note that this holds true even in the presence of non-backbone mediated correlations *à la* eq. (21), since *α*′′ = *µ*(1 + *κ*) *>* 0. The correlations thus vanish as Δ*t* → 0 and produce only a sub-leading contribution to the short time limit considered here.

How does this observation—short time dynamics of the center of mass of *N* discrete loci is independent of the correlations encoded by *κ*—square with the short-time limit for the center of mass of a finite subchain, which manifestly depends on *κ*? To answer this question, note that the boundaries of the short and long time limits for the discrete loci do not match: the short time limit applies for lag times over which individual loci move less than the separation between the closest pair, such that *z*_min_ ≡ min_*i*≠*j*_ *z*_*ij*_ ≫ 1, while the long time limit applies for lag times over which individual loci move more than the separation between the most distant pair, *z*_max_ ≡ max_*ij*_ *z*_*ij*_ ≪ 1. So there exists an intermediate regime *z*_min_ ≲ 1 ≲ *z*_max_, where the behavior is non-universal in general (but can of course

still be predicted from eq. (23)). It is this intermediate regime which will reproduce the previously discussed short time dynamics of the center of mass of a continuous subchain, if we consider the special case of evenly spaced loci, 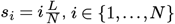, and take *N* → ∞. The short time regime discussed in this section, in contrast, does not exist for the continuous subchain, since in that case 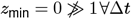 (note that technically here, the minimum in *z*_min_ should be replaced with an infimum; one might also write *z*_min_ = 0^+^).

#### Example: continuous Rouse chain

We can identify the continuous Rouse chain as a specific instance of our generic scale-free model by comparing eq. (17) to the known expression for the MSD of two loci on a continuous Rouse polymer (e.g. eq. (2.88) in [66]). One obtains 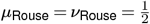 and

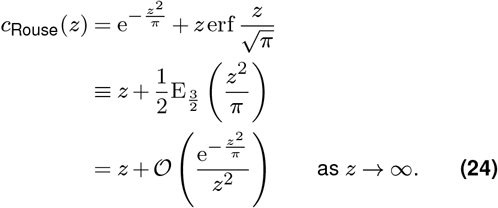

Thus, as *z* → ∞ the approach *c*_Rouse_(*z*) → *z* is faster than polynomial; we write *κ* = ∞ and recover 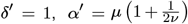. The latter can be rewritten as

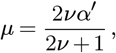

the exponent relation for freely draining chains found in [54]. Our treatment clarifies that this relation holds for any (scale-free) model where short-time displacement correlations are short-ranged (*δ*′′ *>* 1) and generalizes to eq. (10) otherwise.

## Notes

### Competing Interest Statement

The authors have declared no competing interest.

### Summary of Updates

Updated framing of results and discussion of implications/outlook

